# The C-terminal PARP domain of the long ZAP isoform contributes essential effector functions for CpG-directed antiviral activity

**DOI:** 10.1101/2021.06.22.449398

**Authors:** Dorota Kmiec, Maria-José Lista-Brotos, Mattia Ficarelli, Chad M Swanson, Stuart JD Neil

## Abstract

The zinc finger antiviral protein (ZAP) is a broad inhibitor of virus replication. Its best-characterized function is to bind CpG dinucleotides present in viral RNA and, through the recruitment of TRIM25, KHNYN and other cellular RNA degradation machinery, target them for degradation or prevent their translation. ZAP’s activity requires the N-terminal RNA binding domain that selectively binds CpG-containing RNA. However, much less is known about the functional contribution of the remaining domains. Using ZAP-sensitive and ZAP-insensitive human immunodeficiency virus type I (HIV-1), we show that the catalytically inactive poly-ADP-ribose polymerase (PARP) domain of the long ZAP isoform (ZAP-L) is essential for CpG-specific viral restriction. Mutation of a crucial cysteine in the C-terminal CaaX box that mediates S-farnesylation and, to a lesser extent, the inactive catalytic site triad within the PARP domain, disrupted the activity of ZAP-L. Addition of the CaaX box to ZAP-S partly restored antiviral activity, explaining why ZAP-S lacks CpG-dependent antiviral activity despite conservation of the RNA-binding domain. Confocal microscopy confirmed the CaaX motif mediated localization of ZAP-L to vesicular structures and enhanced physical association with intracellular membranes. Importantly, the PARP domain and CaaX box together modulate the interaction between ZAP-L and its cofactors TRIM25 and KHNYN, implying that its proper subcellular localisation is required to establish an antiviral complex. The essential contribution of the PARP domain and CaaX box to ZAP-L’s CpG-directed antiviral activity was further confirmed by inhibition of severe acute respiratory syndrome coronavirus 2 (SARS-CoV-2) replication. Thus, compartmentalization of ZAP-L on intracellular membranes provides an essential effector function in the ZAP-L-mediated antiviral activity.

**Author summary:** Cell-intrinsic antiviral factors, such as the zinc-finger antiviral protein (ZAP), provide a first line of defence against viral pathogens. ZAP acts by selectively binding CpG dinucleotide-rich RNAs, which are more common in some viruses than their vertebrate hosts, leading to their degradation. Here, we show that the ability to target these foreign elements is not only dependent on ZAP’s N-terminal RNA-binding domain, but additional determinants in the central and C-terminal regions also regulate this process. The PARP domain and its associated CaaX box, are crucial for ZAP’s CpG-specific activity and required for optimal binding to cofactors TRIM25 and KHNYN. Furthermore, a CaaX box, known to mediate post-translational modification by a hydrophobic S-farnesyl group, caused re-localization of ZAP from the cytoplasm and increased its association with intracellular membranes. This change in ZAP’s distribution was essential for inhibition of both a ZAP-sensitized HIV-1 and SARS-CoV-2. Our work unveils how the determinants outside the CpG RNA-binding domain assist ZAP’s antiviral activity and highlights the role of S-farnesylation and membrane association in this process.

## Introduction

Cell-intrinsic antiviral factors are an important line of defence against viral pathogens. Although diverse in structure and function, these proteins often share common characteristics including broad antiviral activity conferred by targeting common aspects of viral replication, interferon-stimulated gene expression and rapid evolution due to selective pressures imposed by pathogens [1]. The zinc finger antiviral protein (ZAP) is a broadly active antiviral protein that is induced by both type I and II interferons and is under positive selection in primates [2][3][4][5]. It restricts reverse transcribing viruses, RNA viruses and DNA viruses as well as endogenous retroelements, with retroviruses and positive-strand RNA viruses being the most common viral systems to study ZAP [6].

ZAP’s broad antiviral activity relies on binding viral RNA, thereby either inhibiting their translation and/or target them for degradation by interacting with cellular cofactors such as the 3’-5’ exosome complex, TRIM25, KHNYN and OAS3-RNaseL [7][8][9][10][11][12][13][14][15]. There are four characterized ZAP isoforms, with the long (ZAP-L) and short (ZAP-S) isoforms being the most abundant [3][16]. All ZAP isoforms contain an N-terminal RNA-binding domain (RBD) and a central domain that binds poly(ADP)-ribose [7][17]. However, ZAP-L and ZAP-S differ in that ZAP-L contains a catalytically inactive C-terminal poly (ADP ribose) polymerase (PARP) domain [3]. ZAP distinguishes between self and non-self RNA at least in part by selectively binding CpG dinucleotides [18][19][20]. These are present at a low frequency in vertebrate genomes due to cytosine DNA methylation and spontaneous deamination of the 5-methylcytosine to thymine [21]. Many vertebrate viruses, including RNA viruses that do not have a DNA intermediate, also have a much lower CpG frequency than expected based on the mononucleotide composition of the viral RNAs and this is likely to be due at least in part to restriction by ZAP [22][18][23][24][25].

ZAP was originally identified as a restriction factor for murine leukemia virus and can target several different retroviruses including primary isolates of HIV-1 [7][8][26][27][28]. ZAP more efficiently targets CpGs in the 5’ region of HIV-1 *env* than other regions of the viral genome and introducing CpGs into this region creates a highly ZAP-sensitive HIV-1 [18][14][28]. This model ZAP-sensitive virus has been used to discover and characterize ZAP cofactors such as TRIM25 and KHNYN [14][29]. While the RNA binding domain (RBD) of ZAP is crucial for its selectivity [19][20], much less is known about the functional relevance of the other domains and motifs.

We aimed to determine the functional relevance of ZAP’s domains and their contribution to the mechanism of CpG-specific antiviral activity. In addition to the RBD, we identified that the PARP domain and CaaX box found in ZAP-L, but not ZAP-S, are required for antiviral activity against CpG-enriched HIV-1 and SARS-CoV-2, explaining why ZAP-L is much more antiviral against these viruses than ZAP-S. Both the PARP domain and CaaX box were required for optimal interaction with ZAP cofactors KHNYN and TRIM25. Our findings explain the difference in activity between the two main isoforms of ZAP and highlight the functional contribution of C-terminal regions to the control of important human pathogens such as HIV-1 and SARS-CoV-2.

## Results

### Both the RNA-binding domain and C-terminal domains of ZAP contribute to its CpG specific activity

Full-length ZAP contains an RNA binding domain consisting of four zinc finger domains, a central domain comprised of a fifth CCCH zinc finger, two WWE domains and a C-terminal PARP domain (Fig 1A)[7][10][3][17]. Early studies using rat ZAP suggested that an N-terminal portion of the protein containing the four zinc fingers is sufficient for antiviral activity against murine leukemia virus (MLV) and Sindbis virus [7][30]. However, this was characterised using overexpression experiments in cells expressing endogenous ZAP and the endogenous and exogenous proteins could multimerise [31][32], complicating the experimental interpretation. To compare how much of ZAP’s activity can be attributed to the RBD itself, we initially tested two truncation mutants of the protein, containing either the first 256 amino acids or the last 649 amino acids in ZAP CRISPR KO HEK293T cells. Co-transfection of full-length ZAP with wild type HIV-1 NL4-3 or HIV-1 _*env*86-561_CpG (mutant containing additional 36 CpG dinucleotides introduced into *env* nucleotides 86–561 [14], referred to in this manuscript as HIV-1 CpG-high) resulted in a modest inhibition at the highest concentration. In contrast, a potent dose-dependent inhibition of HIV-1 CpG-high was observed with wild type ZAP (Fig 1B dashed lines). The N-terminal or C-terminal portions of ZAP did not inhibit infectious virus yield, virion production or viral protein expression (Fig 1B right panel and Fig S1A). In line with previous reports [4][31] the phenotype of RBD deletion could be phenocopied by five alanine substitutions in the proposed RNA binding groove (V72/Y108/F144/H176/R189 – 5xRBM) [4] (Fig 1C). Moreover, mutation of residues that directly interact with a CpG dinucleotide, Y108A or F144A, abolished ZAP antiviral activity for HIV-1 CpG-high [19][20]. Of note, these mutations have been reported to relax the CpG-specificity for ZAP antiviral activity on HIV-1 [19]. However, we did not observe antiviral activity for ZAP Y108A or F144A on wild type HIV-1 (Fig 1C-D), which is consistent with the complete loss of function phenotype for these mutations previously observed for Sindbis virus [20]. Thus, the RBD of ZAP is essential yet insufficient for CpG-mediated restriction of HIV-1, implying important effector functions elsewhere in the protein.

**Figure 1.**
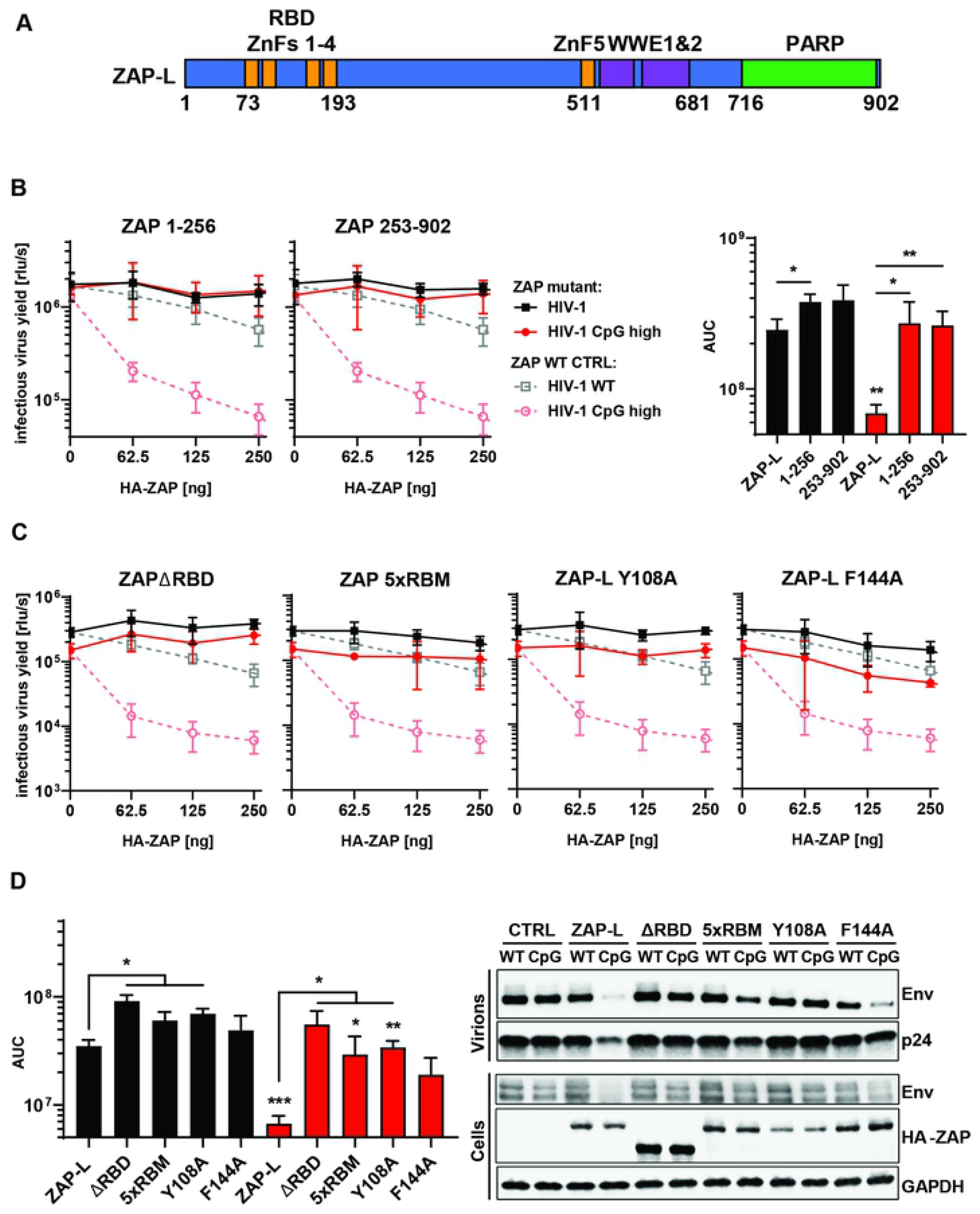
RNA binding is crucial for ZAP’s antiviral activity. (A) Schematic showing domain organisation of long isoform of ZAP (ZAP-L): four N-terminal zinc fingers form RNA binding domain (RBD), fifth zinc finger (ZnF5) and two WWE domains are located in the central part and catalytically inactive Poly(ADP-ribose) polymerase (PARP) domain is at the C-terminal part. (B) Infectious virus yield measured by TZM-bl infectivity assay in relative light units per second [rlu/s] obtained from HEK293T ZAP KO cells co-transfected with wild type (WT; black) HIV-1 and CpG-enriched mutant (CpG-high; red) viruses and increasing doses of pcDNA HA-ZAP constructs encoding the full length ZAP-L (WT CTRL; dashed line), 1-256aa or 253-902aa parts of the protein (solid lines) (left panel). Area Under the Curve (AUC) calculated from the same titration experiments (right panel). (C) Infectious virus yield from HEK293T ZAP KO co-transfected with WT and mutant virus and increasing concentration of pcDNA HA-ZAP with truncated ZAP 253-902 (ΔRBD), ZAP-L mutant unable to bind RNA (V72A/Y108A/F144A/H176A/R189A; 5xRBM) or ZAP-L with substitutions at positions in direct contact with bound RNA CpG (Y108A and F144A) and (D, left panel) derived AUC values. Mean of n=3 +/- SD. Significant differences are indicated as: * p < 0.05; ** p < 0.01. Stars directly above CpG virus values (red bars) indicate statistically significant difference as compared to WT virus (black bar) tested with the same ZAP variant and (right panel) representative western blot of produced virions and ZAP transfected (250ng) cells showing viral Env and Gag (p24) proteins as well as HA-tagged ZAP and GAPDH loading control.

To determine domains required for CpG-specific antiviral activity outside the RBD, we tested ZAP mutants carrying deletions in the central domains (ZnF5 and WWE1 or WWE2) or the PARP domain (Fig 2A). While deletions in the central domain partly reduced ZAP antiviral activity, deletion of the PARP domain resulted in an almost complete loss of CpG-specific inhibition (Fig 2A and B). All PARP proteins except for ZAP (PARP13) and PARP9 can catalyse the transfer of ADP-ribose to target proteins [33]. This lack of catalytic ability has been suggested to be caused by a deviation from the conserved triad motif “HYE” that is required for NAD+ cofactor binding and PARP catalytic activity as well as the partial occlusion of the active site by a salt bridge between H809 and Y824 on one side, and a short alpha helix between residues 803 and 807 at the other [34][35]. Interestingly, residues found to be under strong positive selection in primates (Y793, S804, F805) - often a hallmark of pathogen-host interactions - are located in this alpha helix [3] (Fig 2C). Also, while ZAP orthologs from some tetrapods appear to have an intact catalytic motif similar to PARP12, substitutions in the human ZAP’s PARP domain within the canonical NAD+ binding site prevent the protein from binding this substrate [29][34]. Despite this, mutation of the residues that are present in the triad motif positions, Y786, Y818 and V875 to alanine or H-Y-E abolishes ZAP’s inhibition of Sindbis virus [35], suggesting that the structural integrity of the ZAP PARP domain provides important function.

**Figure 2.**
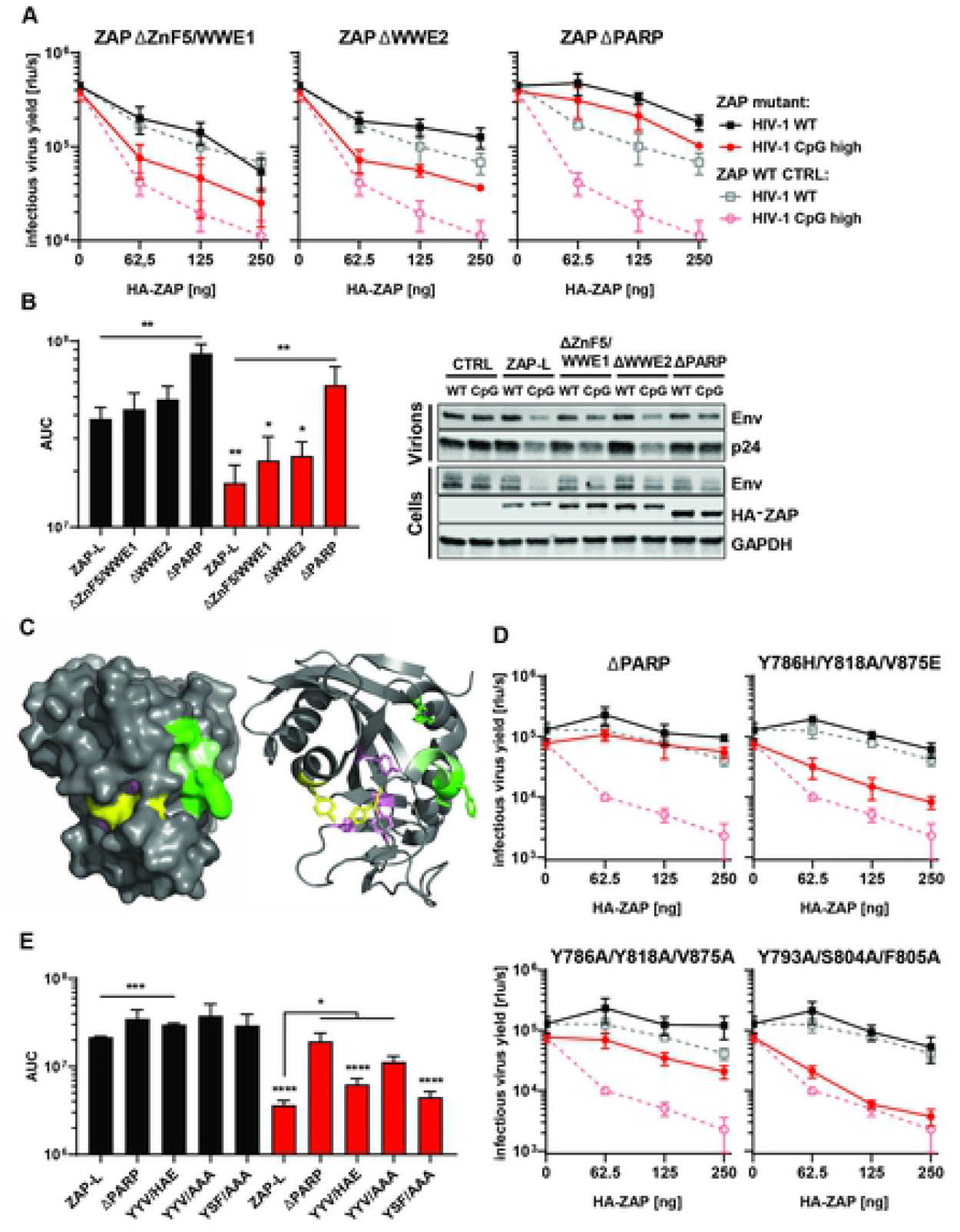
Determinants of ZAP’s function located outside RBD. (A) Infectious virus yield from HEK293T ZAP KO co-transfected with WT (black) and mutant (red) virus and increasing concentration of pcDNA HA-ZAP-L control (WT CTRL, dashed line) or mutated pcDNA HA-ZAP with deleted ZnF5 and first WWE domain (Δ511-563; ΔZnF5/WWE1), second WWE (Δ594-681; ΔWWE2) or PARP domain (Δ716-902; ΔPARP). (B) Corresponding AUC values and representative western blot (250ng). (C) Position of studied residues in crystal structure of ZAP’s PARP domain. Residues under positive selection in primates are shown in green, canonical triad positions in pink and residues forming the salt bridge which closes the NAD+ binding grove are shown in yellow. (D) Infectious virus yield from HEK293T ZAP KO co-transfected with WT (black) and mutant (red) virus and increasing concentration of pcDNA HA-ZAP-L control (WT CTRL, dashed line), missing PARP domain or carrying amino acid substitutions in alternate triad motif (Y786H/Y818A/V875E, Y786A/Y818/V875A), or residues under positive selection (Y793A/S804A/F805A) (solid lines). (E) Corresponding AUC values. Mean of n=3 +/- SD. * p < 0.05; ** p < 0.01; *** p<0.001.

To determine if these residues modulate CpG-specific antiviral activity and explain the apparent lack of inhibition by C-terminally truncated ZAP, we mutated ZAP’s residues 786, 818, 875 (canonical triad positions, pink) and 793, 804 and 805 (sites under positive selection, green) within the PARP domain (Fig 2C). Mutation of the Y786, Y818 and V875 to alanine resulted in a large loss of antiviral function (Fig 2D and 2E) though this was also associated with a substantial decrease in ZAP expression (Fig 2E). Mutation of these residues to H-A-E did not alter ZAP expression but led to a significant loss of antiviral activity. Meanwhile, alanine substitutions at positions under positive selection did not affect the antiviral phenotype (Fig 2D). Therefore, the residues in these positions in ZAP-L that constitute the triad motif in catalytically active PARPs, but not the rapidly evolving residues within the PARP domain, contribute to CpG-specific viral inhibition. However, this does fully account for the loss of phenotype observed with deletion of ZAP’s C-terminus (ΔPARP).

### The C-terminal CaaX box is crucial for ZAP antiviral activity against CpG-enriched HIV-1

While ZAP-L has been reported to be more active than ZAP-S lacking the C-terminal domain (Fig 3A) against at least some viruses, this remains contested and may be virus-specific [3][5][36][37][38][39][40][41]. In agreement with data obtained with the C-terminally truncated mutant ΔPARP (Fig 2), ZAP-S displayed no significant CpG-specific HIV-1 antiviral activity (Fig 3B). We also tested whether co-expression of both isoforms could have synergistic activity and found that ZAP-S had no significant effect on the CpG-high HIV-1 virus even in the presence of ZAP-L (Fig S2).

**Figure 3.**
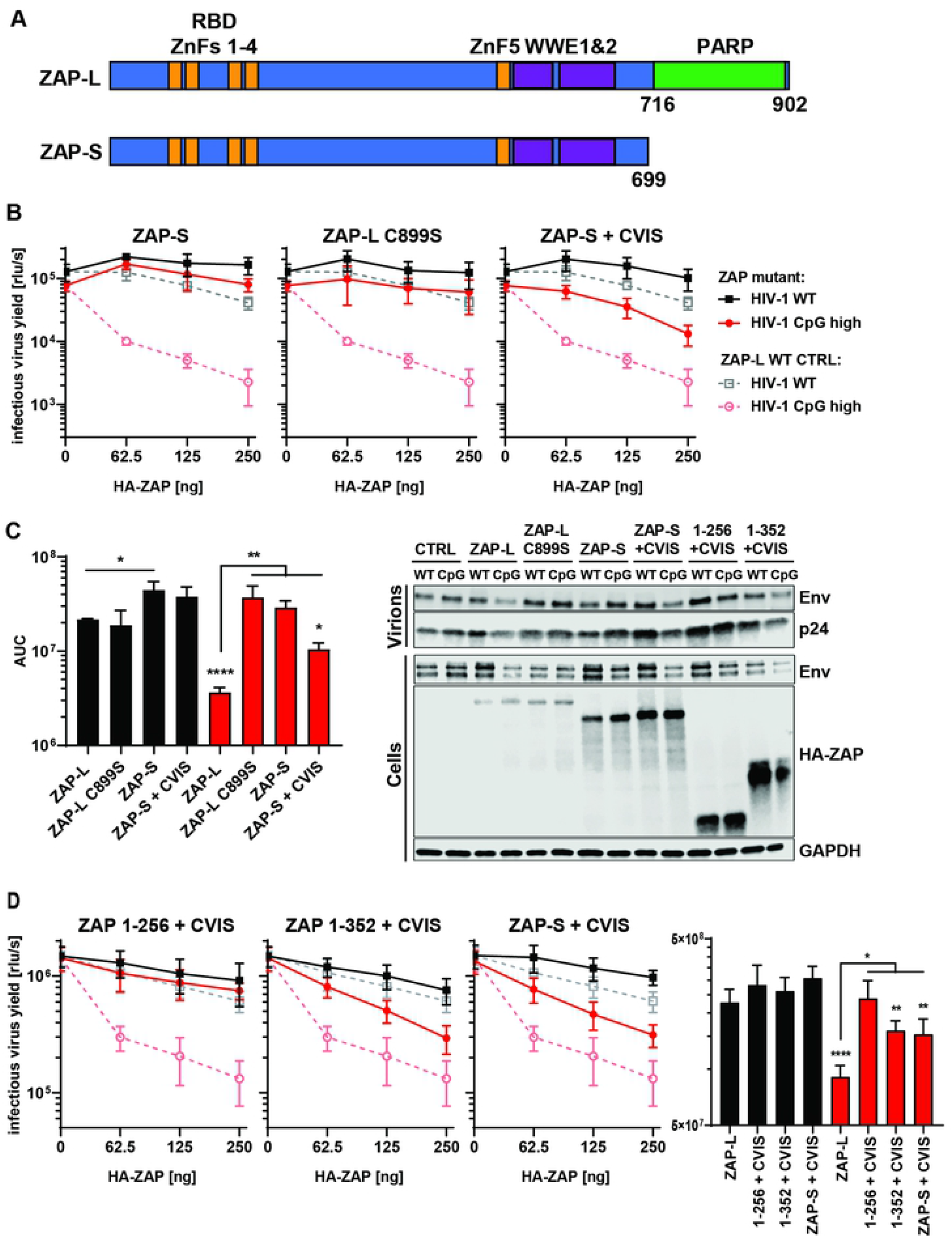
Contribution of CaaX motif to ZAP’s antiviral activity. (A) Schematic showing ZAP-L, which contains PARP domain and CaaX motif (amino acids “CVIS”) and short isoform of ZAP (ZAP-S). (B) Infectious virus yield from HEK293T ZAP KO co-transfected with WT (black) and mutant (red) virus and increasing concentration of ZAP-L CTRL (dashed line), ZAP-S, ZAP-L with mutated crucial cysteine (C899S) in CaaX or ZAP-S with added CVIS motif (solid lines). (C) Corresponding AUC values and representative western blot (250ng). (D) Infectious virus yield from HEK293T ZAP KO co-transfected with both viruses and pcDNA encoding truncated ZAP (1-256 or 1-352) with added CVIS motif and corresponding AUC values. Mean of n=3-5 +/- SD. * p < 0.05; ** p < 0.01; **** p<0.0001.

The ZAP-L PARP domain ends with a well-conserved CVIS sequence that forms a CaaX box (Fig S3B), which mediates a C-terminal post-translational modification through the addition of hydrophobic S-farnesyl group [41]. To evaluate the contribution of the S-farnesylation motif, we mutated the cysteine in the ZAP-L CaaX box to serine (C899S) and added the CaaX box to ZAP-S (ZAP-S + CVIS). The C899S mutation in ZAP-L completely abolished its antiviral activity while addition of the CaaX box to the C-terminus of ZAP-S resulted in a substantial increase in inhibition of HIV-1 CpG-high (Fig 3B-C). Thus, the CaaX box is essential for ZAP antiviral activity on CpG-enriched HIV-1 and can significantly enhance ZAP-S activity even in the absence of the PARP domain. We also analysed whether the N-terminus of ZAP was sufficient for antiviral activity in the presence of the CaaX box. The CVIS motif was added to the first 256 or 352 amino acids of ZAP. The addition of the CVIS motif to ZAP 1-352 led to a partial rescue of CpG-specific activity, comparable to that observed in the case of ZAP-S + CVIS (Fig 3D), though it did not add antiviral activity to ZAP 1-256, suggesting that there might be additional determinants of antiviral function present in the 256-352 region. Because the closest paralogue to ZAP, PARP12, does not share a conserved CpG binding motif or CaaX motif found in mammalian and bird ZAPs (Fig S3A and B), we tested the wild type and modified protein containing CVIS motif (PARP12 + CVIS) by overexpression in ZAP KO HEK293T cells (Fig 4A-D). PARP12 had no antiviral activity against either WT or CpG-enriched HIV-1, and the addition of ZAP’s CaaX or RBD was not sufficient to promote a gain of function phenotype. However, a chimeric PARP12 containing the ZAP RBD in addition to the CaaX box gained partial antiviral phenotype similar to ZAP 1-352 + CVIS, highlighting functional differences between these paralogs in both the RNA binding domain and the C-terminal PARP-domain govern antiviral function.

**Figure 4.**
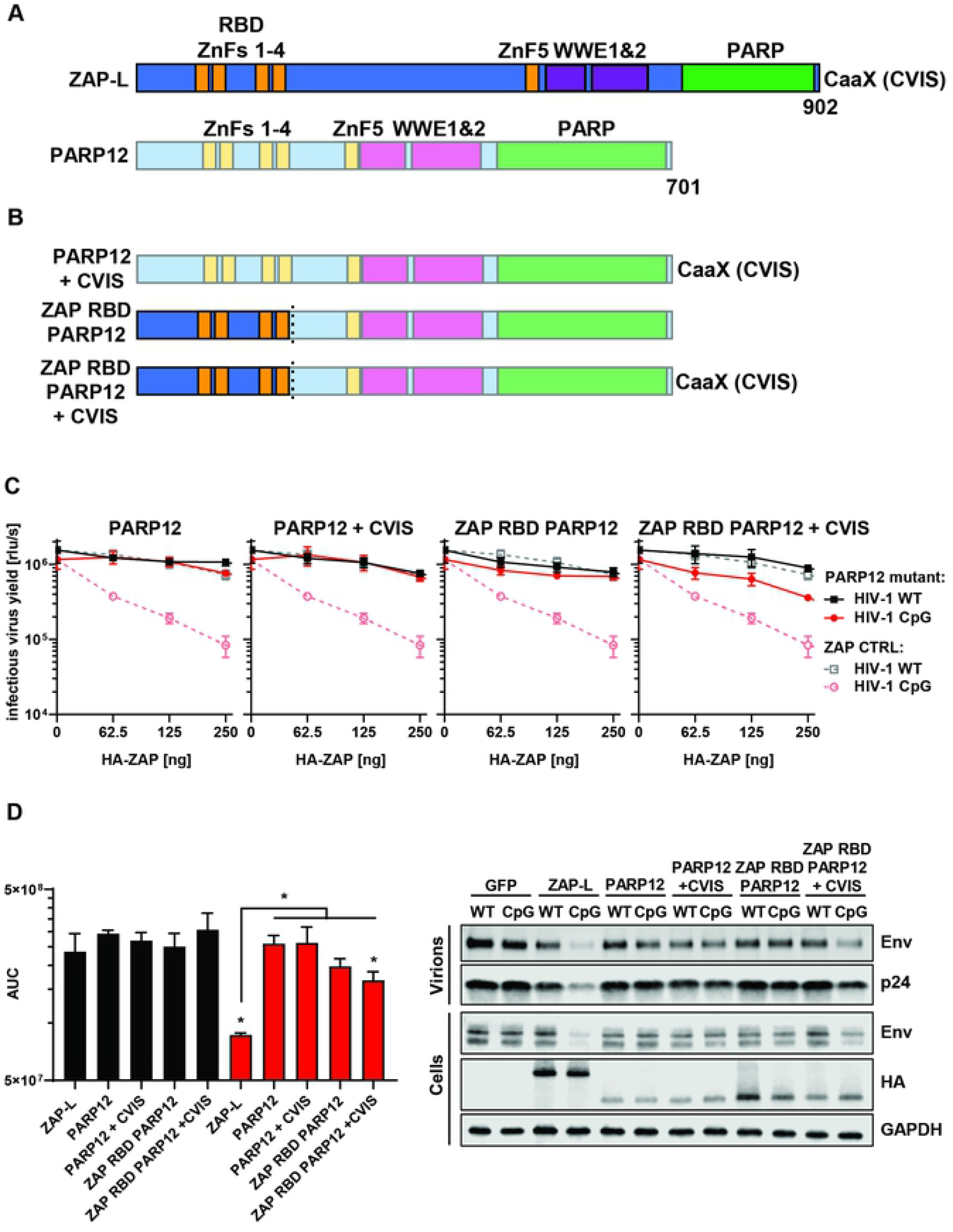
Determinants of CpG-specific antiviral function in ZAP and PARP12. (A) Schematic showing the domain organisation of ZAP-L and PARP12. (B) Schematic of generated PARP12/ZAP chimeric constructs. (C) Infectious virus yield from HEK293T ZAP KO co-transfected with WT (black) and mutant (red) virus and increasing concentration of pcDNA HA-ZAP-L CTRL (dashed line), PARP12, or ZAP/PARP12 chimera (solid lines) and (D) corresponding AUC values and representative western blot (250ng). Mean of n=3+/- SD. * p < 0.05

ZAP-L S-farnesylation has been hypothesized to direct it to endocytic membranes to target incoming viruses that enter cells through endocytic pathways and replicate in viral replication compartments derived from cellular membrane invaginations, such as Sindbis virus [41][42][37]. However, the experiments above tested ZAP antiviral activity on transfected provirus constructs, which effectively start the viral replication cycle at gene expression and bypasses viral entry and the other pre-integration steps. Furthermore, HIV-1 does not replicate in compartments formed from cellular membranes like positive strand RNA viruses. Therefore, the CaaX box cannot be required for ZAP-L to target incoming HIV-1 and intracellular membranes could be used as a platform to establish an antiviral complex. To confirm that ZAP localization to membranes was dependent on the CaaX motif, we generated GFP-tagged versions of wild type and mutant ZAP. Importantly, the GFP-tag did not interfere with ZAP-L antiviral activity (Fig S4). Confocal microscopy of live HEK293T ZAP KO cells transfected with GFP-ZAP (Fig 5A) showed that ZAP-S localized mainly to the cytoplasm, while ZAP-L accumulated in the intracellular vesicular compartments [41][37]. The localization pattern for ZAP-L and ZAP-S was reversed for ZAP-L C899S and ZAP-S + CVIS, respectively. Therefore, vesicular localization appears to correlate with antiviral activity for CpG-enriched HIV-1 (compare Fig 3B and 5A). By contrast, both GFP-ZAP-L and GFP-ZAP-S localized to stress granules defined by G3BP puncta upon poly(I:C) transfection and this was not affected by mutation or transfer of the CaaX box (Fig S5). To determine if the localization observed in the microscopy experiments was also linked to the increased association of ZAP with cellular membranes, we isolated the cytoplasmic (C), membrane (M) and insoluble fractions (D) of HEK293T cells and found that ZAP-L, but not ZAP-S, was present in the membrane enriched fraction. This association could be disrupted by washing the cell lysates in 0.5M salt buffer, while such treatment did not affect membrane association of calnexin (Fig S6), suggesting that ZAP farnesylation mediates only a weak association with the cytoplasmic face of target membranes. Isolation of cytoplasmic and membrane fractions from ZAP-transfected KO HEK293T cells confirmed that while ZAP-L was present at comparable levels in both fractions, the distribution of the ZAP-L C899S mutant resembled that of cytoplasmic ZAP-S, G3BP and GAPDH (Fig 5C). However, while ZAP-S-CVIS relocalizes to resemble ZAP-L localization, its membrane association failed to survive the subcellular fractionation, suggesting a weaker interaction. This, in keeping with its only partial gain of antiviral activity (Fig 3B), further indicates the importance of the integrity of the PARP domain in ZAP-L activity.

**Figure 5.**
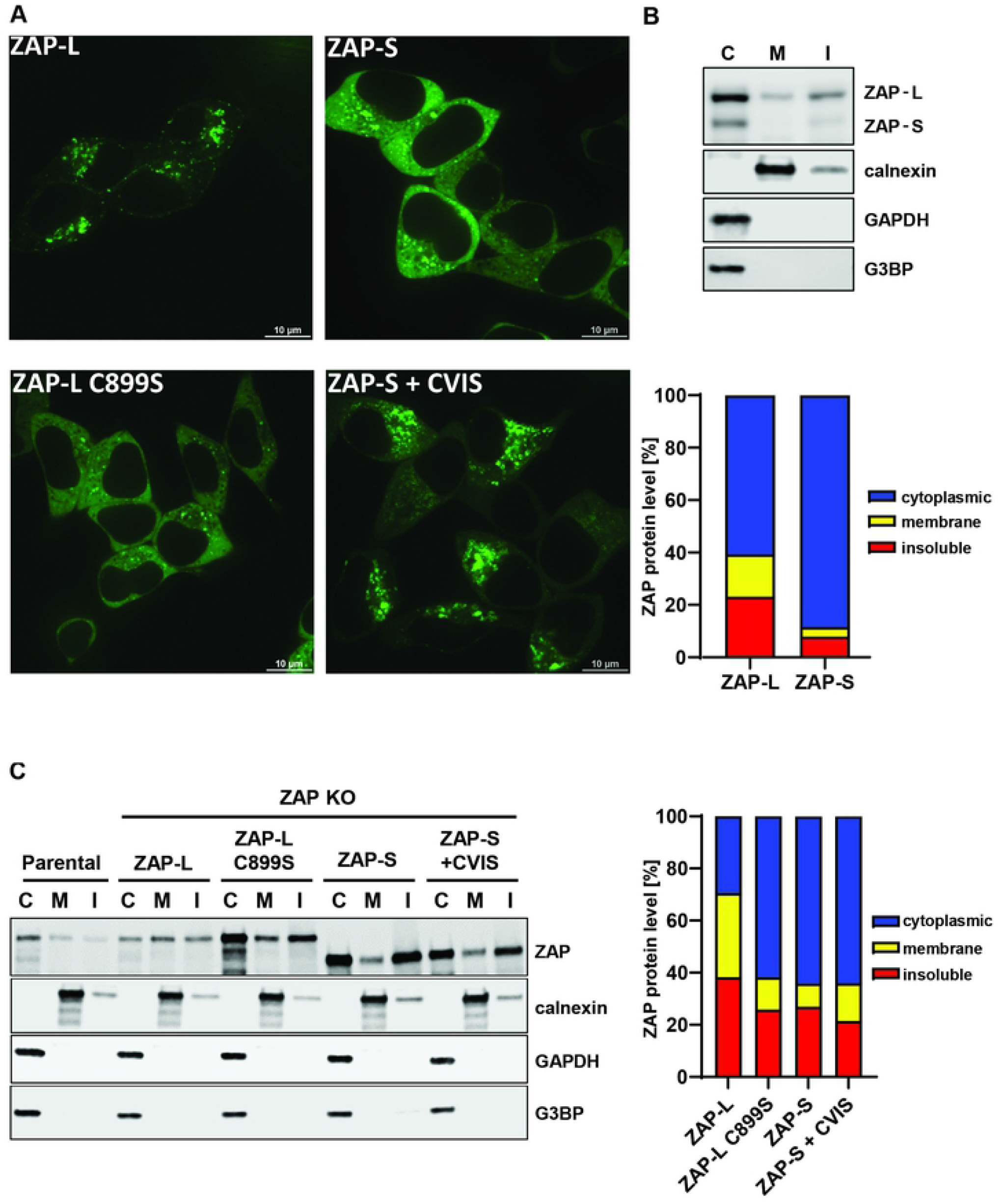
Cellular distribution of ZAP-S, ZAP-L and their CaaX motif mutants. (A) Confocal microscopy images of live HEK293T ZAP KO cells 24h after transfection with 250ng of GFP-tagged ZAP isoforms or ZAP-L with inactivated CaaX (C899S) and ZAP-S with added CaaX motif (+CVIS). Size bar 10μm. (B) Representative western blot and quantification of ZAP present in cell fractionation samples of parental HEK293Ts (mean of n=5) or (C) ZAP KO cells following transfection of 60ng HA-ZAP constructs (mean of n=3). Cytoplasmic (C), membrane (M) and insoluble (I) fractions are shown. Calnexin serves as a marker for membrane protein and G3BP and GAPDH are cytoplasmic protein controls.

We then determined whether ZAP targeting to intracellular membranes is required for its interaction with ZAP cofactors to mediate its antiviral activity against CpG-enriched HIV-1. Pulldown of GFP-tagged ZAP isoforms and mutants revealed that ZAP-L coimmunoprecipitated with endogenous KHNYN more efficiently than ZAP-S, and the 1-256 and 1-352 truncation mutants bound even lower levels of KHNYN (Fig 6A and B). The same pattern was observed for TRIM25. ZAP-S containing the CaaX box showed a gain of interaction with the cofactors. However, even without the functional CaaX box, ZAP-L bound more KHNYN than ZAP-S, indicating that both S-farnesylation, as well as the PARP domain itself, likely play important roles in this interaction.

**Figure 6.**
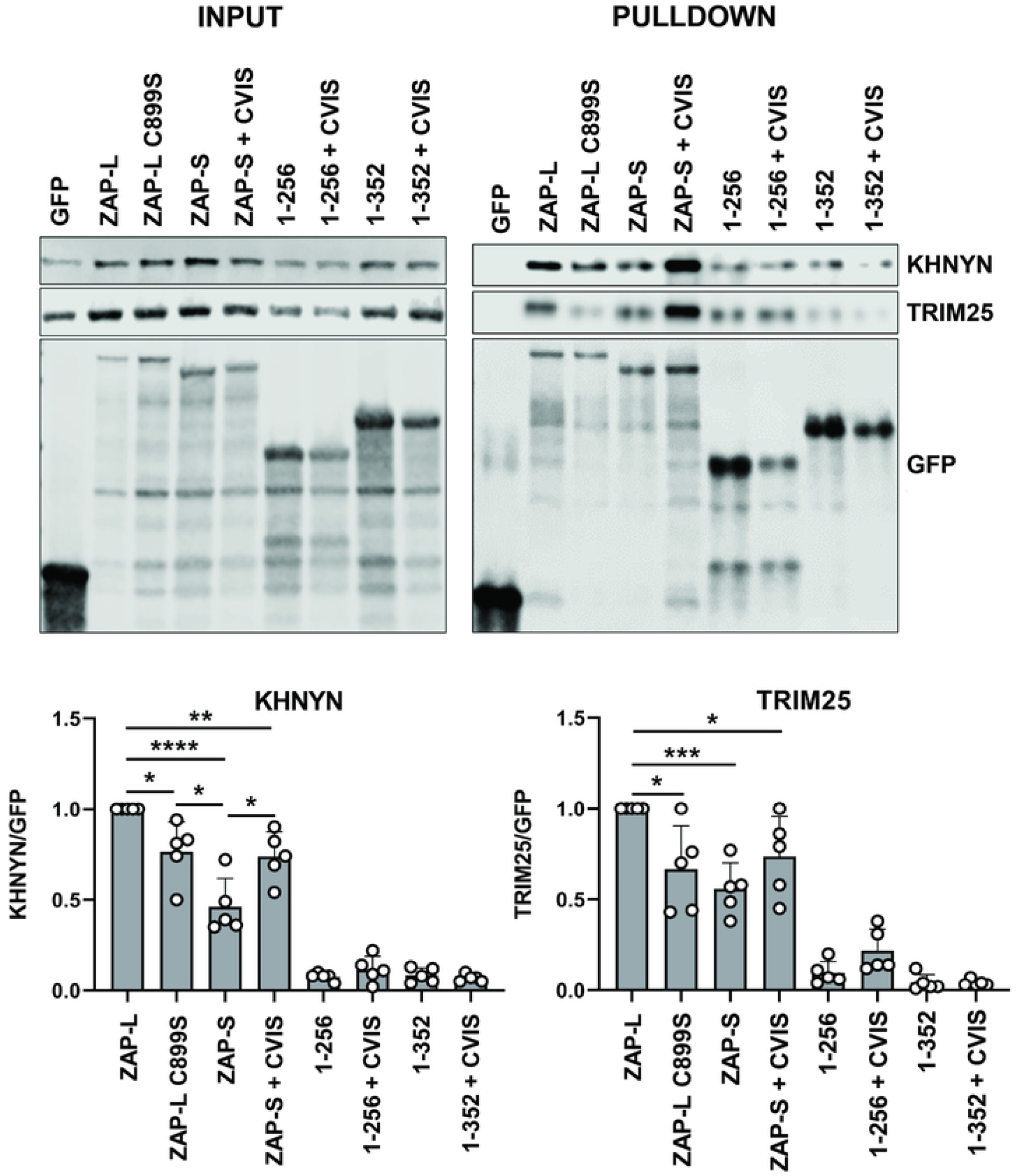
Binding of ZAP to cofactors KHNYN and TRIM25. Upper panel: Representative western blot of GFP-ZAP isoforms and mutants overexpressed in HEK293T ZAP KO cells and co-immunoprecipitated using GFP-binding magnetic beads. Input and pulldown samples were stained for GFP as well as endogenous KHNYN and TRIM25. Lower panel: quantification of pulled-down KHNYN and TRIM25 normalized to relative GFP signal. Mean of n=5 + SD. * p < 0.05; ** p < 0.01; *** p < 0.001; **** p < 0.001.

### The CaaX box and PARP domain are required for ZAP antiviral activity against SARS-CoV-2

Having established determinants of ZAP required to restrict a virus that produces its RNAs in the nucleus, we then sought to confirm these data with an RNA virus that replicates exclusively in the cytoplasm. SARS-CoV-2 has recently been reported to be restricted by ZAP, particularly after exposure of cells to interferon gamma [5], and replicates in double membrane vesicle compartments derived from the ER [43], in contrast to the Sindbis virus replication compartments created by membrane invaginations in the plasma and endosomal membranes [42]. To test if ZAP determinants required to inhibit CpG-enriched HIV-1 also are required for the restriction of SARS-CoV-2, we co-transfected ZAP KO HEK293T cells with plasmids encoding human ACE2 and the indicated ZAP isoform or mutant protein, followed by infection with SARS-CoV-2 at MOI 0.01. Detection of intracellular N protein and viral RNA in the supernatants two days post-infection confirmed that ZAP-S restricts this virus to a far lesser degree than ZAP-L (Fig 7) [5]. ZAP-L restriction was completely abolished when the CaaX box was mutated and transferring this motif to ZAP-S significantly increased its antiviral activity. ZAP-L also required the CpG binding residue Y108 and the YYV motif in place of the PARP catalytic triad for full antiviral activity against SARS-CoV-2. Thus, the determinants of restriction for ZAP-L are similar for a retrovirus that does not replicate on cellular membranes and SARS-CoV-2 which replicates in viral replication compartments derived from the ER.

**Figure 7.**
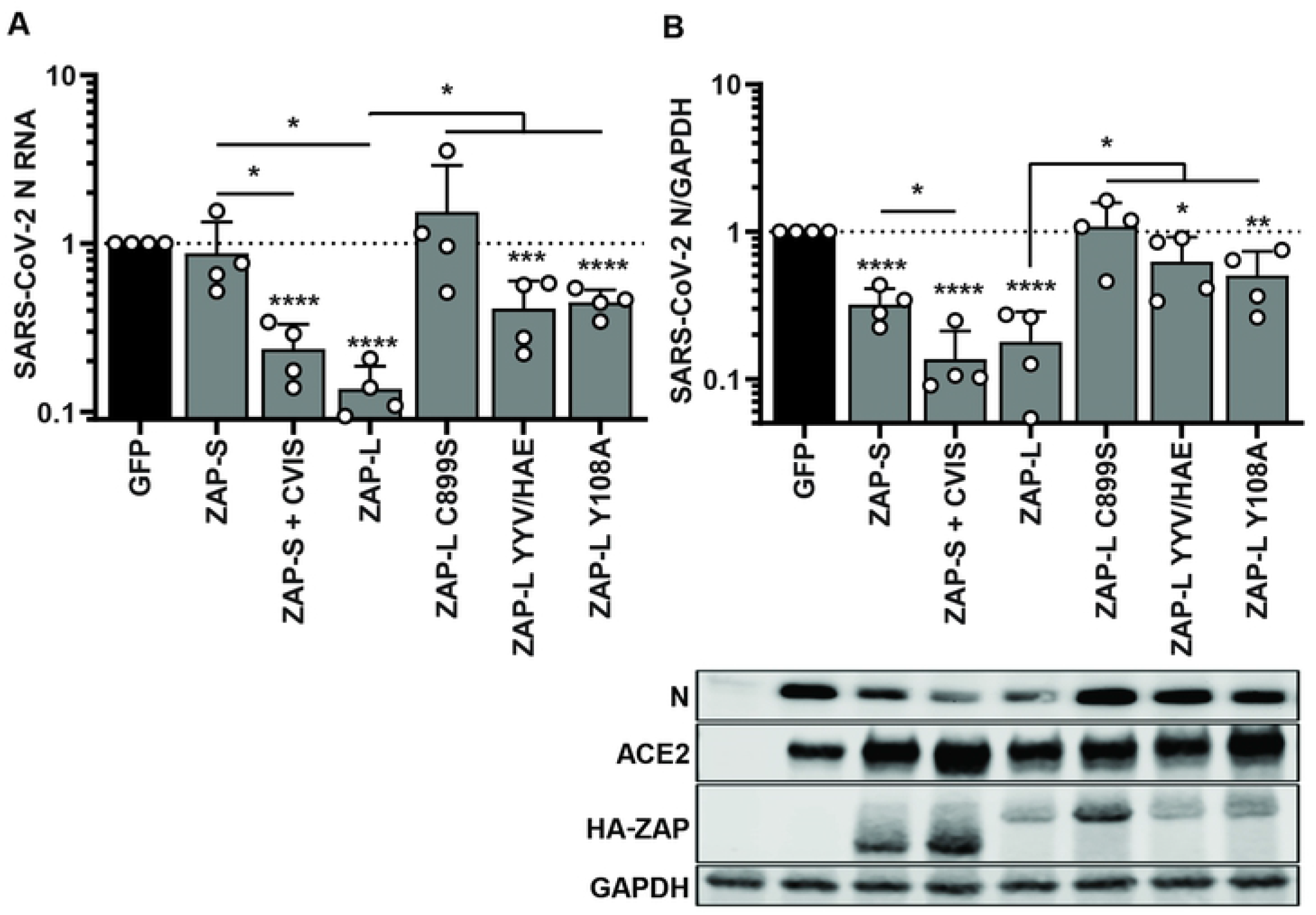
Role of identified ZAP motifs in the inhibition of SARS-CoV-2. (A) Viral RNA produced in HEK293T ZAP KO cells transfected with pcDNA encoding human ACE2 and indicated ZAP isoforms/ mutants or GFP control plasmid, 48h after infection with SARS-CoV-2 England 2 strain at 0.01 MOI. Quantification of qRT-PCR detecting viral nucleocapsid (N) RNA in the cell supernatant and (B) SARS-CoV-2 N levels in the infected cells, with a representative western blot (lower panel). Mean of n=4 + SD. * p < 0.05; ** p < 0.01; *** p < 0.001; **** p < 0.001.

## Discussion

In this study, we demonstrate that in a robust knockout cell-based system ZAP-L, but not ZAP-S, can efficiently inhibit both a CpG enriched HIV-1 as well as SARS CoV-2. We further demonstrate that the C-terminal PARP domain, and particularly its associated farnesylation motif, is essential for this differential activity. Interestingly, while ZAP-L and ZAP-S are derived through alternative splicing and polyadenylation, ZAP-L is expressed constitutively in most cells, whereas ZAP-S expression is more variable and upregulated by type 1 and 2 interferons. Several studies, including this one, have shown that human ZAP-L has more potent antiviral function than ZAP-S for alphaviruses, retroviruses and coronaviruses [3][41][16][37]. However, human ZAP-S clearly has potent antiviral activity for some viruses, such as human cytomegalovirus, when it is the only isoform expressed [44][16][5][45]. Both ZAP-L and ZAP-S have been shown to regulate cellular mRNA expression, so the different isoforms may have differential activity, depending on the transcript [46][47][4][37]. Interestingly, in the context of our experimental system, ZAP-S has no antiviral activity on its own, nor does it augment or interfere with the activity of ZAP-L.

While both ZAP-L and ZAP-S differ only at the C-terminus, their N-terminal RBDs are identical. Within this four zinc finger domain, ZnF2 specifically accommodates a CpG in its binding pocket, and mutation of the contact residues in ZAP-L that define this specificity abolish CpG-dependent restriction of both HIV-1 and SARS CoV-2. This contrasts with a previous study that suggested that such mutations at Y108 and F144 lose their specificity for CpG, but broaden ZAPs antiviral activity to wild-type HIV-1 sequences [19]. The reason for this discrepancy is unclear, although, the previous study did involve the ectopic expression of TRIM25 in a TRIM25/ZAP double knockout cell. Furthermore, as expected from the lack of antiviral activity for ZAP-S, neither the core RBD alone (1-256) or an extended version (1-352) have antiviral activity. Thus, while essential for RNA-binding and CpG specificity, the RBD likely has no intrinsic antiviral activity alone at physiological expression levels. A recent preprint has suggested that ZAP-S can inhibit SARS CoV-2 by negatively regulating the −1 frameshift between Orf1a and Orf1b [48]. Consistent with this, we do see a small reduction in N expression in infected cells expressing ZAP-S alone, but it is insufficient to significantly impact viral production in the supernatant. The Y108A mutant in ZAP-L substantially reduced antiviral activity against SARS CoV-2 indicating that ZAP-L targets SARS CoV-2 via CpG dinucleotides.

The C-terminal PARP domain of ZAP-L is catalytically inactive but ends with a CaaX-box farnesylation motif, CVIS. The CVIS sequence mediated ZAP-L relocalization from the cytoplasm to intracellular membranes. This association appears relatively weak, in line with evidence that protein farnesylation itself is not sufficient for stable association with membranes [49][50]. As such, this may suggest a dynamic exchange of ZAP-L between membrane binding and the cytosol would allow ZAP-L also to localize to cytoplasmic stress granules [4][46]. ZAP-L has been shown to localize to endosomal compartments, but other studies have also indicated that ZAP associates with the ER and nuclear membranes as well [51][41][4][37]. Appending the CaaX box to ZAP-S and even the ZAP 1-352 fragment was sufficient to confer antiviral activity against both against CpG-enriched HIV-1 or SARS CoV-2, in agreement with previous data with Sindbis virus [37]. However, given that the HIV-1 RNA is being targeted after transcription and export from the nucleus, and SARS CoV-2 during exclusively cytoplasmic replication, it is unlikely that ZAP farnesylation is targeting incoming viruses or specific membrane bound replication compartments per se as has been suggested for Sindbis virus, especially considering the differences between alphavirus and coronavirus replication compartments. Rather, farnesylation is more likely to allow compartmentalization or assembly of macromolecular complexes on non-self CpG-rich viral RNAs to facilitate their downstream inactivation irrespective of the subcellular location of viral replication itself. In keeping with this notion, our data indicates the Caax-box modulates the efficiency of interaction with the essential ZAP cofactors TRIM25 and KHNYN. Lipid modification is a common feature of other antiviral proteins including GBP2, GBP5 and the dsRNA sensor OAS1 and is also required for their antiviral function [52][53][54][55]. Moreover, the relocalization of DNA and RNA sensors such as STING and MAVS from the ER or mitochondrial membranes to endolysosomes is coupled to their pattern recognition activities [56]. Importantly, while stress granules have been suggested as a site of ZAP‘s antiviral activity [32], the lack of requirement for the CVIS in this localization argues against their function as a platform for ZAP-L-mediated restriction.

Similar to ZAP’s RBD, the CaaX motif appears to be extremely well conserved in mammals and even birds. A recent study suggested that avian ZAP RBD has lower CpG-specificity than mammalian proteins [29]. It is thus likely that the evolution of CaaX happened after the duplication of genes that gave rise to PARP12 and ZAP, but still preceded the RBD adaptations that enabled efficient CpG-specific viral inhibition. While the CVIS is essential for ZAP-L activity, appending it to ZAP-S or a ZAP-RBD-PARP12 fusion is not sufficient to confer full antiviral activity. This implies that the catalytically inactive PARP, in conjunction with the ZnF5 and WWE domains, plays an important role in ZAP-L function. The two WWE domains and ZnF5 have been a subject of recent pre-print showing that these two regions combine into a single integrated domain that binds ADP-ribose, which facilitates antiviral activity[17]. Furthermore, ZAP was also shown to be mono-ADP-ribosylated by PARP14 and PARP7 [57][58]. Therefore, ZAP is potentially a target for ADP-ribosylation by multiple PARP proteins which can regulate its activity, but it cannot perform this function on its own due to mutations within its PARP domain. We found that residues forming what would be the catalytic triad motif in active PARP domains contribute to ZAP’s antiviral function. It is tempting to speculate that the evolution of CpG-specific antiviral activity enhanced by the PARP domain in ZAP led to, or was a consequence of, the loss of its own ADP ribosylation ability. Despite the inactive catalytic site being occluded in the ZAP PARP domain, mutation of the residues that would form the active site modulates ZAP-L activity and stability, suggesting structural integrity of the PARP domain may facilitate cofactor interactions and/or multimeric assembly on target RNAs. Validation of such hypotheses awaits a full structure of ZAP rather than its constituent domains.

In summary, we show that ZAP-L localization to membranes and the integrity of its C-terminal PARP domain facilitate cofactor recruitment provide an essential antiviral effector function in the context of its ability to bind CpG dinucleotides in viral RNAs.

## Materials and Methods

### Expression constructs and cloning

Previously described pcDNA3.1 HA-ZAP-L and ZAP-S constructs [24] were rendered CRISPR-resistant by introducing synonymous mutations within exon 6. Primers were synthesized by Eurofins, and all PCRs were performed with Q5 High Fidelity DNA Polymerase (NEB). Monomeric enhanced GFP fused to N-terminus of ZAP via a flexible linker (GGGGSGGGGSGGGG) was synthesized by Genewiz and the full-length ZAP cDNA was reconstituted using an internal PsiI site. Specific mutations and deletions were generated using Q5 site-directed mutagenesis or Gibson Assembly (NEB) cloning. pcDNA3.1 HA-PARP12 was generated by PCR amplifying the PARP12 coding sequence (Dharmacon) and ligating into EcoRI/EcoRV sites of pcDNA3.1 using T4 DNA ligase (NEB). Construct sequence identity was confirmed by restriction enzyme digestion and Sanger sequencing (Genewiz). pHIV-1_NL4-3_ and pHIV-1_*env*86-561_CpG were described before [14][23]. pcDNA N-terminally C9-tagged human ACE2 construct was kindly provided by Dr Nigel Temperton.

### Cell lines and culture

Human Embryonic Kidney (HEK) 293T cells were obtained from the American Type Culture Collection (ATCC). Hela and HEK293T CRISPR ZAP KO (exon 6) cells were described previously [14][24]. TZM-bl reporter cells (kindly provided by Drs Kappes and Wu and Tranzyme Inc. through the NIH AIDS Reagent Program) express CD4, CCR5 and CXCR4 and contain the β-galactosidase genes under the control of the HIV-1 promoter [59][60]. Cells were cultured in Dulbecco’s modified Eagle medium with GlutaMAX (Gibco) supplemented with 10% fetal calf serum (FCS), 100 U/ml penicillin and 100 μg/ml streptomycin, and grown at 37°C in a humidified atmosphere with 5% CO2.

### Transfection and HIV-1 infectivity assay

HEK293T ZAP KO cells (0.15-0.2mln) were seeded in 24-well plates and transfected the following day using PEI MAX (3:1 PEI to DNA ratio; Polysciences) with 500 ng pHIV-1 and 0-250 ng pcDNA3.1 protein expression construct. The total amount of DNA was normalized to 1 μg using pcDNA3.1 GFP vector. Media was changed the following day and cell-free virus-containing supernatants and cells were harvested two days post-transfection. To measure infectious virus yield, 10.000/well TZM-bl cells were seeded in a 96-well plate and infected in triplicate. Two days later, viral infectivity was determined using the Gal-Screen kit (Applied Biosystems) according to manufacturer’s instructions. β-galactosidase activity was quantified as relative light units per second using a microplate luminometer.

### SARS-CoV-2 infection

HEK293T ZAP KO cells (0.2 mln) were seeded in 12-well plates. The following day, the cells were transfected using PEI MAX with 100 ng pcDNA C9-ACE2 and either 400 ng pcDNA ZAP or GFP control vector. At 24 hours post-transfection, the cells were infected with SARS-CoV-2 England 2 virus strain at MOI 0.01 (prepared and tested as previously described in [61][62]. After 1 hour (h), cells were washed in PBS to remove the inoculum. Virus-containing cell-free supernatants and cell lysates were harvested two days later.

### Quantitative Real-Time PCR

RNA from infected cell supernatants was extracted using QIAamp viral RNA mini kit (Qiagen) and cDNA was synthesized using the High Capacity cDNA RT kit (Thermo) following the manufacturer’s instructions. The relative quantity of nucleocapsid (N) RNA was measured using a SARS-CoV-2 (2019-nCoV) CDC qPCR N1 and control RNAseP probe set (IDT DNA Technologies). qPCR reactions were performed in duplicates with Taqman Universal PCR mix (Thermo) using the Applied Biosystems 7500 real-time PCR system. Relative SARS-CoV-2 RNA amounts were calculated using the ΔΔCt method.

### SDS-PAGE and immunoblotting

HIV-1 virions were concentrated by centrifugation at 18,000 RCF through a 20% sucrose cushion for 1.5 hours at 4°C. Cells were lysed in radioimmunoprecipitation assay (RIPA) buffer containing cOmplete EDTA-free protease inhibitor (Roche) and 10U/ml benzonase nuclease (Santa Cruz). Cell lysates and concentrated virions were then reduced in Laemmli buffer and boiled for 10min at 95°C. Samples were separated on gradient 8-16% Mini-Protean TGX precast gels (Bio-Rad) and transferred onto 0.45 μm pore nitrocellulose. Membranes were blocked in 5% milk and probed with mouse anti-HA (#901514, Biologend), rabbit anti-HA (#C29F4, Cell Signalling), rabbit anti-GAPDH (#AC027, Abclonal), mouse anti-G3BP (#611126, BD), rabbit anti-calnexin (#ab22595, abcam), rabbit anti-ZAP (#GTX120134, GeneTex), rabbit anti-GFP (#ab290, abcam), mouse anti-KHNYN (#sc-514168, SantaCruz), mouse anti-TRIM25 (#610570, BD), rabbit anti-SARS-CoV-2 N (#GTX135357, GeneTex), rabbit anti-ACE2 (#ab108209, abcam), mouse anti-HIV-1 p24 [63] or rabbit anti-HIV-1 Env (#ADP20421, CFAR), followed by secondary DyLight conjugated anti-mouse 800 (#5257S, Cell Signalling), anti-rabbit 680 (5366S, Cell Signalling), HRP conjugated anti-mouse (#7076S, Cell Signalling) or anti-rabbit (#7074S, Cell Signalling). HRP chemiluminescence was developed using ECL Prime Reagent (Amersham). Blots were visualized using LI-COR and ImageQuant LAS 4000 Imagers.

### Co-immunoprecipitation

HEK293T ZAP KO cells were seeded at 0.3-0.4 mln/ml in 10 cm dishes and transfected the following day with 10 μg pcDNA GFP or pcDNA GFP-ZAP plasmid using PEI MAX. Cells were harvested two days later and ZAP was immunoprecipitated using GFP-Trap magnetic agarose kit (Chromotek) following the manufacturer’s instructions.

### Confocal microscopy

For live-cell microscopy, ~75.000 HEK293T ZAP KO cells were seeded onto poly-Lysine coated 24-well glass-bottom plates and transfected with 250 ng pcDNA3.1 GFP-ZAP using PEI MAX. Cells were visualized 24 h later using a 100x oil-immersion objective equipped Nikon Eclipse Ti-E inverted CSU-X1 spinning disk confocal microscope.

To visualize ZAP relocalization to stress-granules, ~50.000 Hela ZAP KO cells were seeded onto poly-Lysine coated 24-well glass-bottom plates and transfected with 125 ng pcDNA encoding GFP-ZAP using LT1 transfection reagent. 40 h post-transfection, cells were transfected with 100 ng poly(I:C) using Lipofectamine 2000 (Invitrogen) and fixed 6 h later in 2% PFA. Cells were blocked and permeabilized for 30min in PBS containing 0.1% TritonX and 5% Normal Donkey Serum (Abcam), stained overnight with mouse anti-G3BP (BD, #611126, 1:200 dilution), followed by 2 h staining with secondary donkey anti-mouse Alexa Fluor 546 antibody (Invitrogen, A10036, 1:500 dilution) and 1μg/ml DAPI.

### Cell fractionation

HEK293T and HEK293T ZAP KO cells (0.6-0.8 mln) were seeded in 6-well plates. The following day, ZAP KO cells were co-transfected using PEI MAX with 60 ng pcDNA HA-ZAP constructs and 940 ng pcDNA3.1 empty vector. Cells were harvested two days later, washed in PBS and processed using ProteoExtract Native Membrane Protein Extraction Kit (Sigma). Soluble cytoplasmic, membrane protein and insoluble fractions were isolated according to the manufacturer’s instructions, with the addition of three 1 ml PBS or high salt washes between extraction buffer I and II. The insoluble debris fraction was resuspended in RIPA buffer, sonicated and reduced in Laemmli buffer by boiling at 95°C for 10min.

### ZAP sequence analysis

Protein sequences of human ZAP-L orthologs were downloaded from the NCBI database (https://www.ncbi.nlm.nih.gov/). Sequences were aligned using ClustalW2 (https://www.ebi.ac.uk/Tools/msa/clustalw2/) and logo plots were generated using WebLogo online tool (https://weblogo.berkeley.edu/logo.cgi) (ref).

### Data analysis

The area under the curve (AUC) and statistical significance (unpaired two-tailed Student’s t-test) were calculated using Prism Graph Pad. Data are represented as mean ± SD.

## Acknowledgments

We thank other members of the Neil and Swanson laboratories for helpful discussions as well as Dr Monica Agromayor and Prof. Juan Martin-Serrano and their group members for advice and assistance with confocal microscopy. We thank Nigel Temperton for generously providing reagents. The following reagents were obtained through the NIH AIDS Research and Reference Reagent Program, Division of AIDS, NIAID, NIH: TZM-bl from Dr John C Kappes, Dr Xiaoyun Wu and Tranzyme Inc; HIV-1 p24 Hybridoma (183-H12-5C) from Dr Bruce Chesebro and Dr Hardy Chen. The Antiserum to HIV-1 gp120 #20 (ARP421) was obtained from the NIBSC Centre for AIDS Reagents. This work was funded by a Deutsche Forschungsgemeinschaft (German Research Foundation) fellowship to DK (Project number: KM 5/1-1), Wellcome Trust Senior Research Fellowship (WT098049AIA) to SJDN, and Medical Research Council grant MR/S000844/1 to SJDN and CMS. This UK funded award is part of the EDCTP2 programme supported by the European Union. MF is supported by the UK Medical Research Council (MR/R50225X/1) and is a King’s College London member of the MRC Doctoral Training Partnership in Biomedical Sciences. The funders had no role in study design, data collection and analysis, decision to publish, or preparation of the manuscript.

## Supporting information

**Figure S1. Effect of expressed ZAP mutants on viral protein levels.** Representative western blots of experiment shown in (A) Fig.1B and (B) Fig.2D.

**Figure S2. Antiviral effect of ZAP-L and ZAP-S co-overexpression.** Infectious virus yield from HEK293T ZAP KO cells co-transfected with wild type (WT; black) HIV-1 and CpG-enriched mutant (CpG-high; red) viruses and increasing doses of pcDNA HA-ZAP constructs encoding ZAP-L (dashed lines), ZAP-S or 1:1 ratio of both isoforms up to 250ng each (solid lines). Values were normalized to infectivity in the absence of ZAP for each virus (100%). Mean of n=5 +/- SD. Lower panel: representative western blot (250ng HA-ZAP).

**Figure S3. Alignment of RBD of ZAP and PARP12 and conservation of ZAP-L’s CVIS motif in mammals and birds.** (A) Alignment of RNA-binding domains of human ZAP and its paralogue PARP12. Four zinc fingers (grey boxes) and ZAP residues interacting with CpG dinucleotide in bound RNA (highlighted in pink) are indicated. (B) Logo plot of C-termini of mammalian and bird ZAP-L orthologues from NCBI database. Highly conserved C-terminal serine (S901) determines targeting by cellular farnesyl transferase which prenylates highly conserved cysteine (C899).

**Figure S4. CpG-specific antiviral activity of HA and GFP tagged ZAP isoforms.** Infectious virus yield from HEK293T ZAP KO cells co-transfected with wild type (WT; black) HIV-1 and CpG-enriched mutant (CpG-high; red) viruses and increasing doses of pcDNA ZAP with N-terminal hemagglutinin tag (HA) or monomeric enhanced green fluorescent protein (GFP) tag. Mean of n=3 +/- SD.

**Figure S5. Re-localization of ZAP isoforms and their CVIS mutants to stress-granules.** HeLa ZAP KO cells were transfected with 125ng GFP-ZAP (green) and stained for stress-granule marker G3BP (red) following treatment with 100ng of poly(I:C). DAPI staining shows cell nuclei (blue).

**Figure S6. Effect of 0.15M-1M NaCl washes on ZAP’s membrane localization.** Western blot and protein quantification following fractionation of HEK293T cells. Cytoplasmic (C), membrane (M) and insoluble (I) fractions are shown, with relative levels of endogenous ZAP-L and ZAP-S, as well as controls calnexin (membrane fraction control), and G3BP, GAPDH and TRIM25 (cytoplasmic fraction controls).

